# Mode III tear resistance of *Bombyx mori* silk cocoons

**DOI:** 10.1101/2023.11.16.567364

**Authors:** Ateeq Ur Rehman, Vasileios Koutsos, Parvez Alam

## Abstract

This paper concerns the tear properties and behaviour of *Bombyx mori* silk cocoons. The tear resistance of cocoon layers is found to increase progressively from the innermost layer to the outermost layer. Importantly, the increase in tear strength correlates with increased porosity, which itself affects fibre mobility. We propose a microstructural mechanism for tear failure, which begins with fibre stretching and sliding, leading to fibre piling, and eventuating in fibre fracture. The direction of fracture is then deemed to be a function of the orientation of piled fibres, which is influenced by the presence of junctions where fibres cross at different angles and which may then acts as nucleating sites for fibre piling. The interfaces between cocoon wall layers in *Bombyx mori* cocoon walls account for 38% of the total wall tear strength. When comparing the tear energies and densities of *Bombyx mori* cocoon walls against other materials, we find that the *Bombyx mori* cocoon walls exhibit a balanced trade-off between tear resistance and lightweightness.

## INTRODUCTION

*Bombyx mori* (*B. mori*) is one of the most useful domesticated species of mulberry silkworms^1, 2^, spinning a silk cocoon to protect itself as it develops morphologically from its pupal stage. Silk fibres have been used in textiles for more than 4000 years^3^, with the earliest example of a woven silk fabric originating in 3630BC^4^. In a typical silk production process, the cocoon is degummed to separate sericin proteins (the glue) from fibroin proteins (the structural thread). Silk fibres are strong yet soft to the touch, flexible, durable^5^, and absorbent to a wide variety of dyes^1^.

During cocoon construction, *B*.*mori* extrudes silk fibres through the spinneret in its mouth^6^. Similar to silk egg cases^7–13^, the *B. mori* cocoon has a protective function^14–19^. It is a sophisticated multilayer non-woven composite comprised of continuous silk fibres connected by a sericin gum-like coating that acts as a microstructural level glue for the fibres^20^. *B. mori* cocoon size range from 30 – 35 mm^2^. A typical cocoon has been reported as having a meridional diameter of 31.57 *±*0.19 mm and an equatorial diameter of 19.01 *±*0.17 mm^21^. Its wall is typically 0.30-0.59 mm thick^14,19,20,22,23^, it has a density between 377-499 kg/m^3 19,20^, and contains 70-80% silk fibroin with the remainder being sericin protein^3,24^. Silk fibre has a triangular or irregular cross-section with an approximated diameter between 16-26*μm*. A single fibre consists of two fibroins, each with a diameter of 7-14*μm*, which are conjoined by a thin sericin layer 2-4*μm* in thickness^14,23,25^.

The cocoon wall of *B. mori* can be divided into 5 to 15 distinct layers, each varying in their of sericin to fibroin compositional ratio, and each exhibiting different microstructures. The innermost layers are comprised of fibres with mean diameters of ca. 16 *μ*m and the number of fibres/mm^2^ is ca. 21. Contrarily, the outermost layers have higher mean fibre diameters of ca. 26 *μ*m but there are only 8 fibres/mm^2^ in these layers^14^. Comparatively, the inner layers of *B. mori* cocoons comprise lower fractions of sericin, but stronger levels of inter-fibre bonding^22,26,27^. The density of *B. mori* layers has been reported as decreasing from the innermost to the outermost layer^28^. It has additionally been noted that porosities between the inner and outer layers differ significantly and are reported in the ranges of 0.42-0.70 and 0.58-0.84, respectively^29^. The density of silk fibre ranges between 1350-1365 kg/m^3 28^.

Tensile strength, tensile modulus, and the toughness of the cocoon wall are of the order: 16.6 – 54 MPa, 300 – 586 MPa, and 1.1 MJm^−3^, respectively^22,23,30,31^. The wall will strain 18 *±*2 % at maximum tensile strength and the breaking strain is typically in the order of 13-35 %^22,31^. Additionally, the elliptical design of silk cocoons has been understood as being of significant benefit to impact damage tolerance, as an ellipsoid can elastically deform on impact in such a way that energy is stored in the ellipsoid and released at a critical level proportional to the impact force^32^.

The hierarchical structure of *B. mori* silk cocoon impacts the mechanical properties of the subdivided layers^19,33,34^. The Young’s modulus, tensile strength, storage modulus, and loss modulus are all reported as being higher in the inner pelade layer (near the pupa) than they are in the thickness-averaged values of the complete cocoon^35^. The strength and modulus of individual layers rises as fibre areal density increases, and as porosity and fibre diameter decreases from the outermost to the innermost layers^23^. The specific modulus and specific strength of the innermost layer have been reported as being the highest at 24 GPa mm^−1^ and 938 MPa mm^−1^, respectively, while they are significantly lower in the outermost layer, at 1.6 GPa mm^−1^ and 159.4 MPa mm^−1^, respectively. Contrarily, strain at peak stress is the lowest in the innermost layer (7.9 %) and highest at the outermost layer (21.3%). Each cocoon layer contributes to its mechanical properties and behaviour, and the layers are held together mostly by sericin as well as by a few cross-linking fibres. As such, interlayer bonding within a cocoon wall is significantly weaker than intralayer bonding^22^.

The cocoon wall is also permeable to moisture and moisture flux in the outer layers is higher than in the inner layers, as the inner layers are generally lower in porosity and are of higher tortuosity^29^. In addition, the cocoon wall is extremely permeable to gases under normal conditions, yet it has the capacity for CO_2_ gating and thermoregulation under more extreme environmental conditions^36,37^, and is thus able to maintain its internal temperature and CO_2_ levels. It is therefore clear that one of the benefits of the distinctive design of the cocoon wall is to offer protection to the resident pupa, whether that be mechanical, gaseous, or thermal^14^.

While the structure-property relationships of a variety of silk fibres have been researched extensively^25,38–43^, knowledge of silk cocoons is more sparsely documented. As mentioned, cocoons are structures that provide protection to pupae and herein we hypothesise that the geometrically complex, hierarchical, and layered structure of a *B. mori* cocoon should provide a certain level of tolerance to tearing. Multiple reports document the tear resistance of silk sheets and silk derived/inspired composites^44–48^, yet as far as we are aware, there has been no work conducted to date on the tear strength of cocoon walls and their layers. Knowledge of this property will enable a more detailed understanding of how cocoons serve to mechanically protect their pupae. As such, this paper aims to fill this knowledge gap and we aim to detail the tear strength of *Bombyx mori* silk cocoons, in relation to their structures.

## MATERIALS AND METHODS

### Tear Testing of Full Cocoon Walls

A modified ASTM D624-00^49^ trouser tear test method was used to enable testing of the smaller-than-standard-sized samples (due to restrictions imposed by the sizes of the *B. mori* cocoons). Cocoon macro measurements were made using a Vernier caliper and thickness measurements were made using a digital screw gauge. Thirty rectangular trouser-tear samples were prepared (n = 30) from thirty individual cocoons by cutting a 20 mm slit in the equatorial direction of the cocoon wall Figure 1(a). The open cocoon walls were tear tested using an Instron 3369 (High Wycombe, UK) with a 1KN load cell. The initial gauge length was 10 mm and testing was conducted at a displacement rate of 10 mm/min. Each of the split halves of the individual sample (i.e. the trouser legs) was held in the Instron grip and was aligned with the centre line of the sample. The uncut end of the specimen was kept free, and this is the part that would experience a tear at a right angle to the line of the force application. Figure 1 (b) provides a schematic of the trouser tear test, and Figure 1(c) shows the experimental setup.

**Figure 1:**
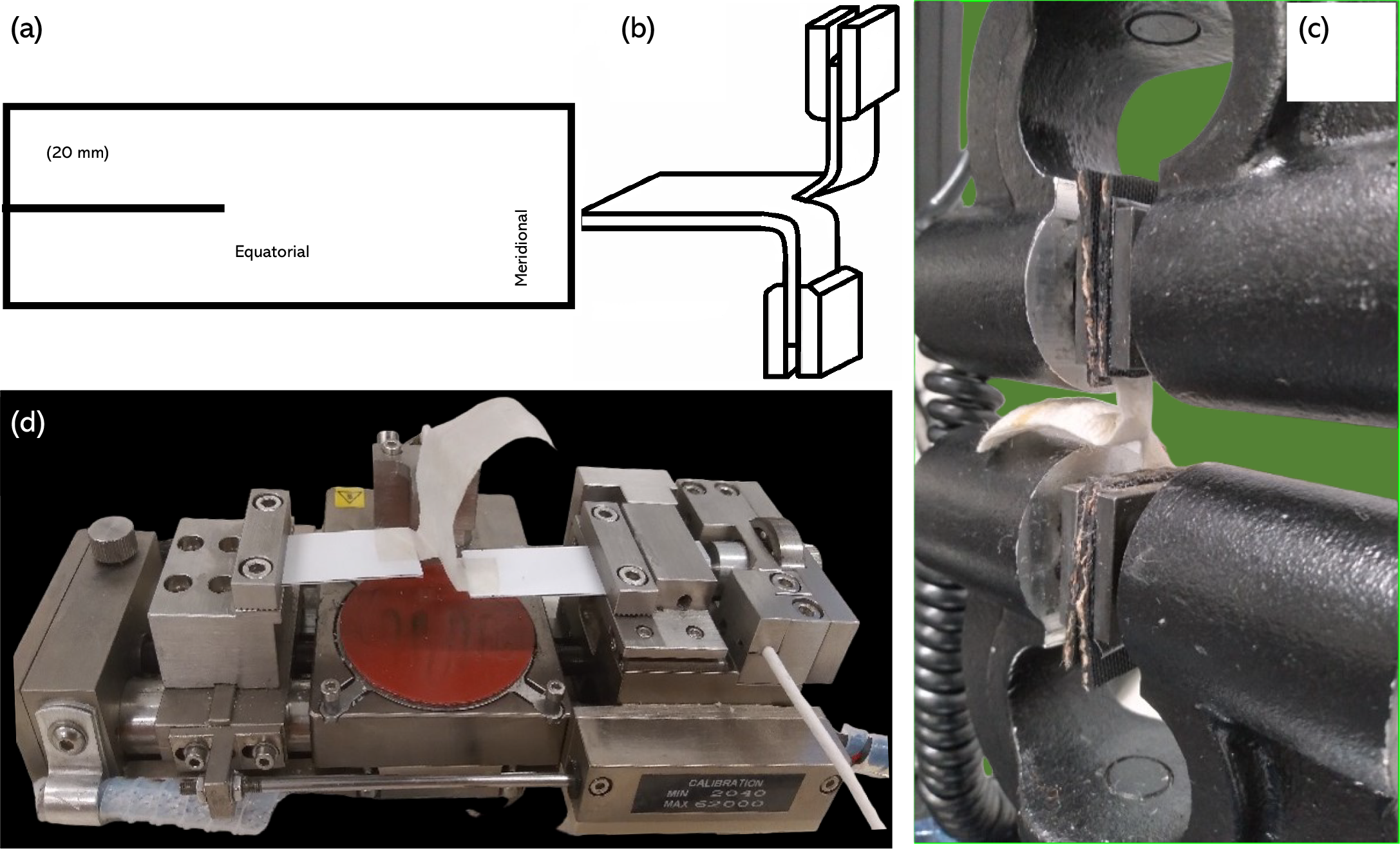
(a) Trouser test full cocoon specimen (b) Trouser Tearing (c) Instron 3369 experimental setup (d) Deben microtensile tester experimental setup.

### Tear Testing of Individual Cocoon Layers

Cocoon walls were separated into seven individual layers. Although individual cocoon walls can in principle be separated into up to fifteen layers, these layers are extremely thin and difficult to test mechanically. As such, we employed a similar approach to Chen et al.^22^ who subdivided the cocoon into seven layers to enable layer-by-layer tensile testing. The seven individual layers were essentially therefore comprised of adjacent layers within the cocoon wall. These cocoon layers (hereinafter: individual layers) were then cut into 20mm wide strips. At the end of each sample, a 10 mm long slit was made in the equatorial direction at the centre fold line of the sample. These were then tested using a Deben (Deben, Suffolk, UK) micro tensile tester with a 5N load cell at an extension rate of 1.5 mm/min. Load was applied to the trouser test specimen and tear force was recorded over a maximum machine extension limit of 11 mm. Figure 1(d) shows a representative sample in example of the experimental setup. The tear strength of the full cocoon walls as well as cocoon layers was calculated using Equation (1) following ASTM D624^49^. Where *S*_*tear*_ is the tear strength, *F*_*tear*_ is the tear force, and *t* is the median thickness of the cocoon wall.

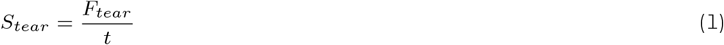

The work done in tearing was calculated using Equation (2). Where *W*_*tear*_ is the work done in tearing, *F*_*tear*_ is the tearing force, and Δ*c* is the tearing length.

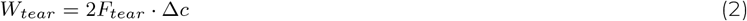

The tearing energy was calculated using Equation (3). Here, *E*_*tear*_ is the tearing energy, *F*_*tear*_ is the tearing force, and *t* is the median thickness of the cocoon wall^50^.

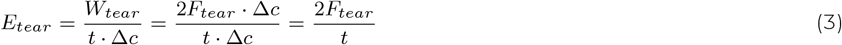

Relative density, *ρ*_*r*_, and porosity, *ϕ*, were calculated using Equation (4) and Equation (5), respectively. Here, *ρ* is the apparent density of the cocoon wall calculated using the formula 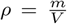, (where *m* is mass and *V* is volume), and *ρ*_*s*_ is the density of *B. mori* silk fibre^51^.

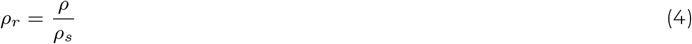

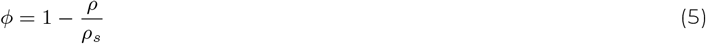

### Digital Microscopy and Image Analysis

To develop our microstructural understanding of tear damage, individual cocoon layers were optically examined using a Dino-Lite digital AM4115-FUT microscope (New Taipei City, Taiwan). This microscope possesses a 1.3-megapixel image capture capability, offering a versatile magnification range from 20*×* to 220*×*. In this paper, images were captured at magnifications up to 60*×*. Customised LEDs emitting UV light in the 375 nm spectrum were used to illuminate the samples. To ensure optimal picture quality and precise control over image acquisition, the microscope was securely affixed to a Dino-Lite RK-10 pedestal as it ensured stable positioning during image capture.

## RESULTS

### General Observations

By degumming in accordance with the procedure described in^31^, the weight percentage of fibroin in native cocoons was determined to be 74% with a standard deviation (SD) of 1.73%. Table 1 presents the physical measurements of thirty cocoons (n = 30) in both meridional and equatorial directions. Cocoon walls conditioned under standard temperature (21 °) and humidity (55%) were used to determine apparent density in Table 1.

**Table 1:** *Bombyx mori* silk cocoon: physical measurements (including the range and the mean *±* the standard deviation (n = 30)

|  | Cocoon Diameter |  | Cut Sheet Dimensions |  | Sheet Thickness | Apparent Density of Wall |
| --- | --- | --- | --- | --- | --- | --- |
|  | Meridional | Equatorial | Meridional | Equatorial |  |  |
|  | (mm) | (mm) | (mm) | (mm) | (mm) | (kg/m<sup>3</sup>) |
| **Range** | 29.3-36.9 | 18.6-23.0 | 15-22.6 | 52.0-63.0 | 0.49-0.92 | 297-389 |
| **Mean** | 32.6 | 20.2 | 19.4 | 56.2 | 0.71 | 331 |
|  | (± 2.0) | (± 1.0) | (± 1.7) | (± 2.3) | (± 0.1) | (± 34) |

### Tear Properties of Cocoon Walls

Tear force (*F*_*tear*_) was determined in accordance with ASTM D624^49^. This standard computes the highest tear force value from the range of available standards, including: BS2782-3 method 360B^53^, BS ISO 34-1:2022^54^, BS EN ISO 6383-1:2015^55^, ASTM D1938-19^56^, ASTM D2261^57^ and BS EN ISO 13937-2:2000^58^. Further details on these standards and comparison curves are provided as Electronic Supplementary Material (Comparison of standard tear force calculations). Using the tear force values (*F*_*tear*_ = 17.5 *±*4.2 N) from 30 samples (cf. Electronic Supplementary Figure S1), where the thickness (*t* = 0.71 *±*0.12 mm), calculations were made to quantify tear strength (*S*_*tear*_ = 25 *±*6.1 kN/m), work done in tearing (*W*_*tear*_ = 9.3 *±*5.5 kN·m) and tear energy (*E*_*tear*_ = 50.1 *±*12.2 kN/m). Two generic tearing characteristics were noted during testing and these were split into groups 1 (G1) and 2 (G2) and characteristics were equally split in the sample set such that 15 of 30 samples showed G1 characteristics, while the other 15 samples exhibited G2 characteristics. Figure 2(a) shows the pictures of representative torn cocoon walls from G1 and G2 and provides additional schematics to represent the generic tear propagation orientations for each group. G1 cocoon walls tearing was generally oriented in the direction of loading, while the tearing of G2 cocoon walls showed an acute angular orientation to the direction of loading. Of the 15 samples exhibiting G2 characteristics of tear propagation, the angles of orientation were found to range from 25 ° (minimum) to 58 ° (maximum), with a mean at 41 ° ±14^*°*^ (SD). Representative tear stress plots are shown for each of the two groups against their tear length, Figure 2(b). In the G1 plot, a gradual decrease can be observed beyond the maximum tear strength. Dissimilarly, a sharper decline in tear stress can be observed beyond the maximum tearing strength in G2 cocoon walls subjected to trouser tearing.

**Figure 2:**
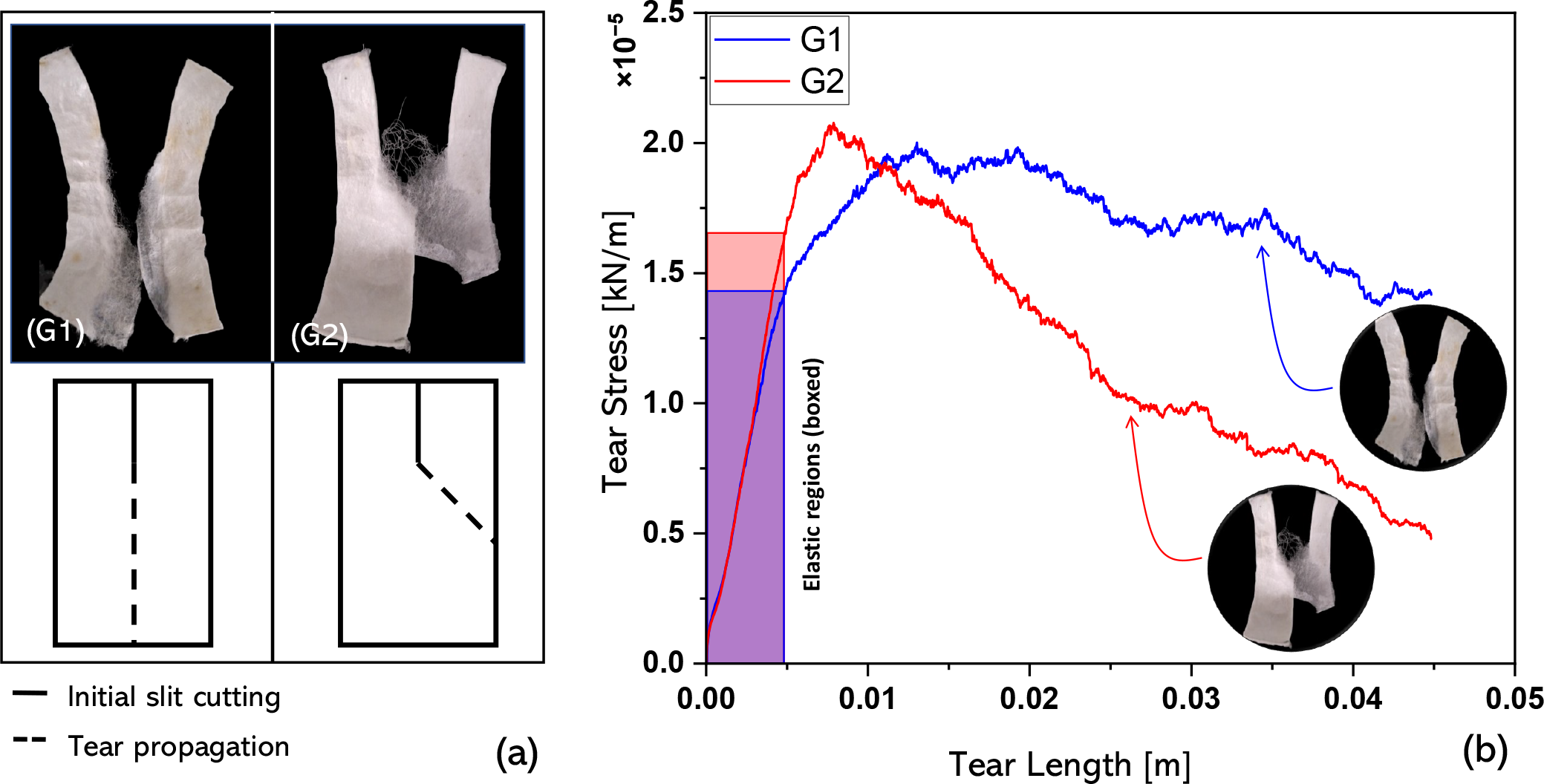
Representative curves from a sample set of 30 cocoons showing tear stress against the tear length for groups 1 (G1) and 2 (G2) generalised tearing orientations. G1 and G2 tearing orientations are shown more clearly on the right hand side of the figure.

To assess whether there are any statistical differences between the two groups in terms of their properties, a one-way ANOVA was conducted with significance levels (*α*) set at 0.001, 0.01, and 0.05. Table 2 provides the results from the one-way ANOVA at *α*=0.05. Since the calculated F-statistic, F = 0.42, is lower than the F critical value of 4.2 (the value against which F is compared), and the P-Value of 0.52 is above *α*, it can be concluded that there is no difference between the mean values of G1 and G2 at the highest selected *α*. The results for *α* = 0.01 and 0.001 were no different, except for that that F critical increases to 7.6 and 13.5 respectively, further reducing the probability of a type 1 error.

**Table 2:** One-way ANOVA between groups 1 and 2 comparing *S*_*tear*_, F is the F-statistic, SS is the sum of squares, MS is the mean sum of squares, d is the degrees of freedom, and F crit is the F-critical value.

| Source of Variation | SS | d | MS | F | P-Value | F crit |
| --- | --- | --- | --- | --- | --- | --- |
| Between the Groups | 15.7 | 1 | 15.7 | 0.42 | 0.52 | 4.2 |
| Within Groups | 1056.8 | 28 | 37.7 |  |  |  |
| Total | 1072.5 | 29 |  |  |  |  |

Additional comparisons between G1 and G2 properties are provided in Table 3, showing the standard deviations (SD) for each sample set as well as the coefficient of variation (CoV). While it can be noted that the average values of G1 for *t, F, S*_*tear*_ and *E*_*tear*_ are within one standard deviation from the arithmetic mean of G2 values for *t, F*_*tear*_, *S*_*tear*_ and *E*_*tear*_, and *vice versa, W*_*tear*_ for G1 is within two standard deviations from the arithmetic mean of *W*_*tear*_ of G2, and *vice versa*. Unlike *E*_*tear*_, the work done (*W*_*tear*_) is a function of tear length and as such, the comparative distance for tearing is higher in G1 than in G2, a higher value of *W*_*tear*_ is expected in the G1 samples.

**Table 3:**
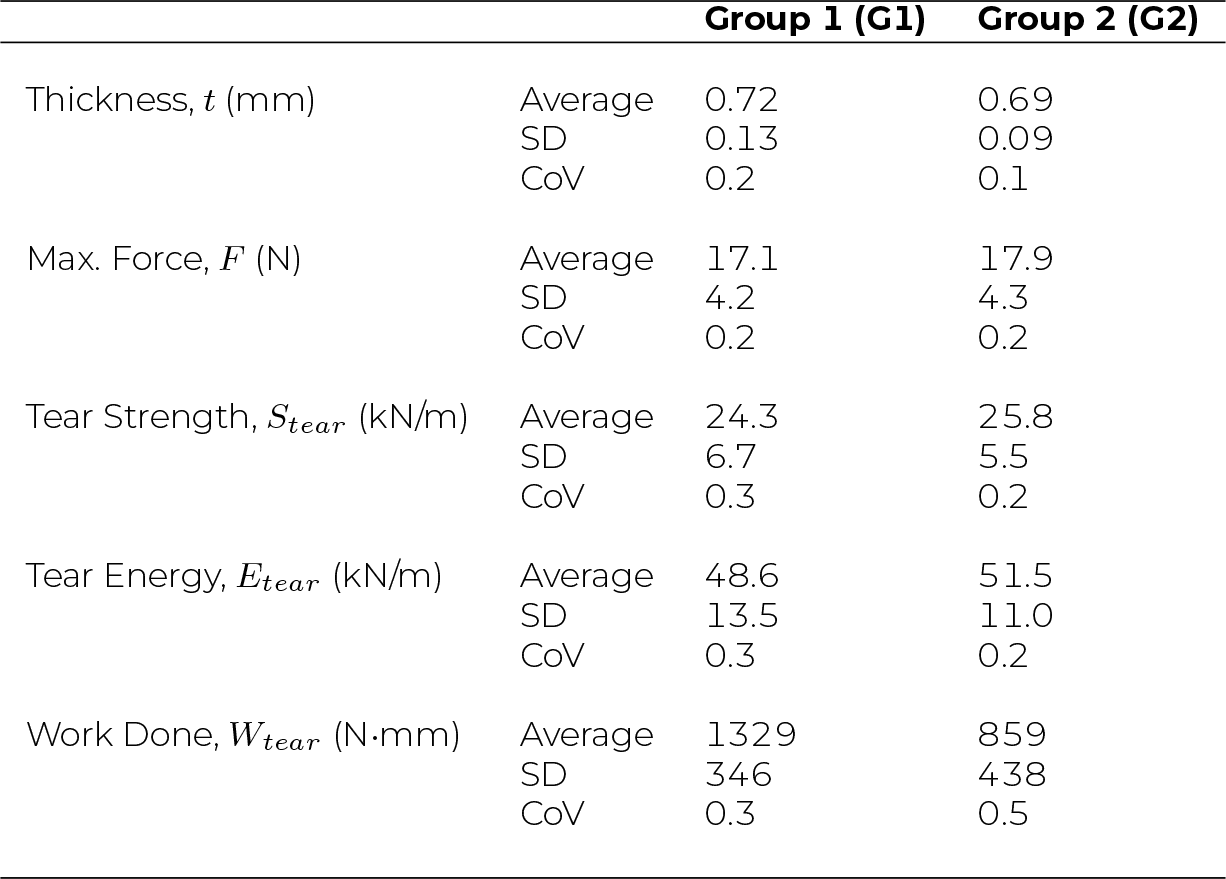
Comparison of tearing parameters among four groups with different tearing behaviours.

A mechanism for *B. mori* cocoon tearing can be suggested, using images of the specimens torn at different stages, Figure 3(a-c). These provide evidence on how tearing both effects, and is affected by, fibre architecture. We find there are three primary fibre failures at the microstructural level that influence the overall mechanism of tearing. In the first Figure 3(a), the applied tearing force initiates both fibre stretching and fibre sliding in the direction of the tear. In the second stage Figure 3(b), we note that fibre stretching and sliding leads into localised fibre piling, a phenomenon previously reported in mechanically loaded *N. cruentata* spider silk egg cases^12^. Here we suggest that similarly to^12^, cross fibres adhered to stretching/sliding fibres get trapped as they slide into other cross fibres/junctions, creating fibre piles. As fibres pile, the applied tearing force required to cause them to deform, displace or break is likely to increase as mechanical energy stores within these reduced mobility fibres. These fibres are thus susceptible to fracture rather than displacement under loading as the tearing energy increases. Tearing will therefore propagate at either a different orientation angle to the axis of tear loading Figure 3(c), or in the axis of loading, depending on the force vectors (magnitude and direction of force), which in turn are influenced by the local cocoon fibre architectures, Figure 3(d-e) and the way in which they experience piling. Given that tearing is a function of deformation in the vicinity of the tear tip^50^, a deviation in deformation from the direction of loading could therefore be associated with the fibre architecture within the cocoon wall. Silkworms build cocoons by overlaying a continuous strand of fibre. As a consequence of this, both larger (major) and smaller (minor) fibre junctions form where fibres are overlaid at different angles. This results in a variable areal density (number of fibres per unit area) as shown in Figure 3 (d). We suggest that there may be some influence to piling at junctions comprising a higher density of fibre cross-overs where there are large numbers of fibres crossing at approximately the same angular orientation, as these may act as nucleation points of some form. Fibre junctions with large numbers of fibres crossing one another would presumably have a greater chance of encouraging the redirection of tearing energies after piling through the nucleation of fibres at these point, and resulting in tear line reorientation. Contrarily, where there is a more uniform distribution of fibres with fewer and smaller fibre junctions are perhaps more likely to permit parallelised piling and may encourage tearing in the axis of loading, Figure 3 (e).

**Figure 3:**
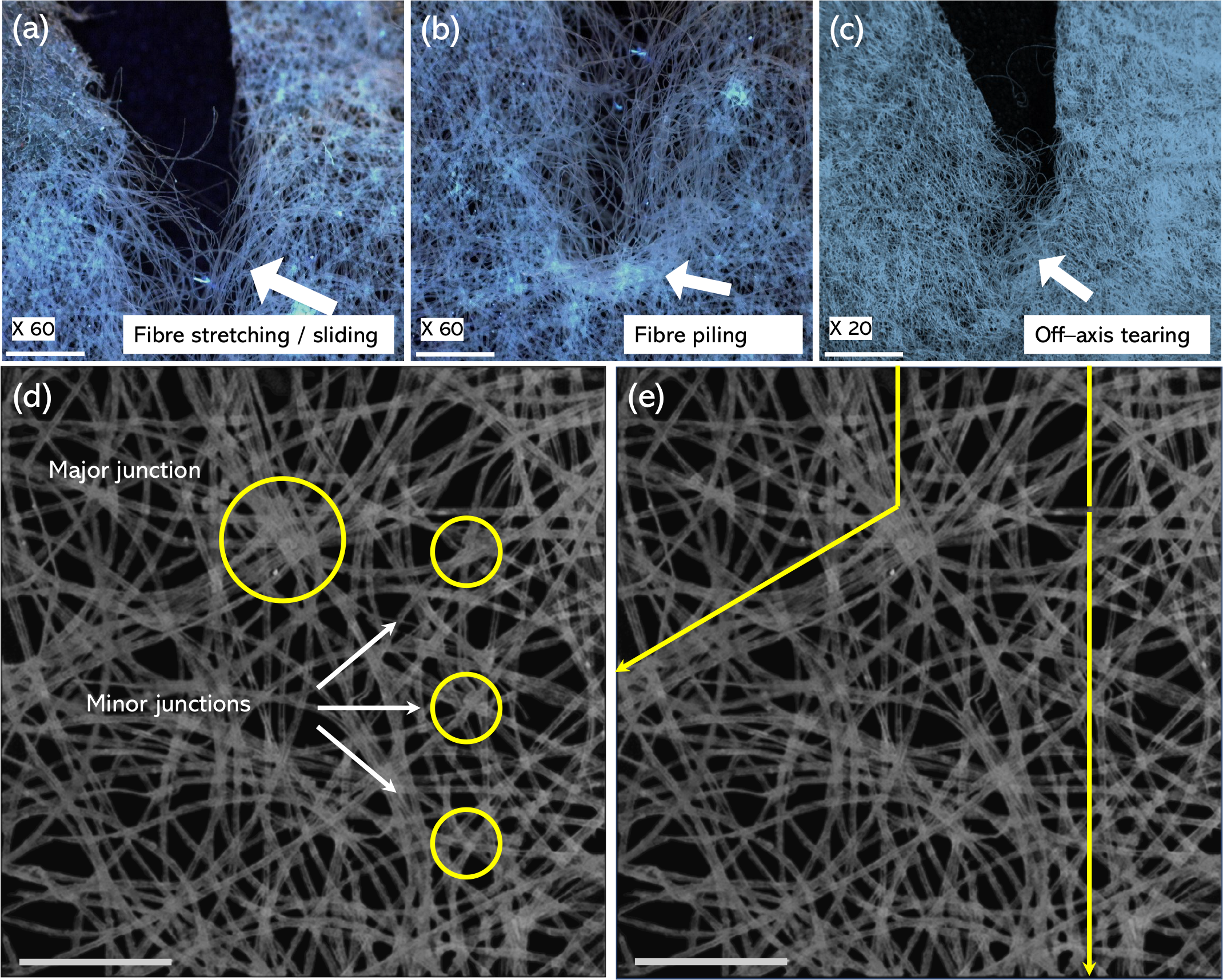
The mechanism of *B. mori* tearing initiates through (a) fibre stretching and sliding followed by (b) the co-movement of cross fibres adhered to the stretching/sliding fibres resulting in local fibre piling and eventuating in (c) the build up of strain energy at fibre piles leading to fracture. Examples of major and minor junctions where fibres cross from layer to layer are shown in (d), while a hypothetical suggestion on how fibre junctions may play a role as nucleation points for differently oriented fibre piling leading to redistributed and reoriented tear energies is shown in (e).

### Tear Properties of Individual Cocoon Wall Layers

Figure 4(a) illustrates the seven subdivided layers of a *B. mori* cocoon, numbered from one (innermost) to seven (outermost). When comparing the properties of the sum of all cocoon layers against the properties of the cocoon wall, we note that both the tear force (*F*_*tear*_), Figure 4(b), and the tear strength (*S*_*tear*_), Figure 4(c), for the complete cocoon wall surpass the cumulative values of the individually tested layers. The mean *F*_*tear*_ of the cocoon wall was measured as 6.6 N higher than the mean sum of *F*_*tear*_ values for the individual cocoon layers. Similarly, there is a ca. 10kN/m *S*_*tear*_ improvement of the complete cocoon wall over the sum of individual layers. The additional resistance to loading in complete cocoon walls is likely attributable to the action of sericin adhering the interfaces of individual layers. The separation of the cocoon into individual layers result in the absence of the additional surface fibre linkages, thus reducing the mechanical properties. Taking the 6.6 N difference therefore as being equally distributed over 6 interfaces in a 7-layer cocoon, we can approximate a 1.1 N overall additional resistance to tearing, per interface. This value exceeds previously reported interlayer peeling loads of 0.32 N^22^ in *B. mori* cocoons, though it should be noted that the earlier reported peeling load were determined using smaller specimens (20 × 5mm), distinct experimental speeds, tensile loading, and different mechanical test machines. Furthermore, the present study is specifically focussed on tear loading, which localises any peeling (and thus interfacial effects) to the area near the tear tip.

**Figure 4:**
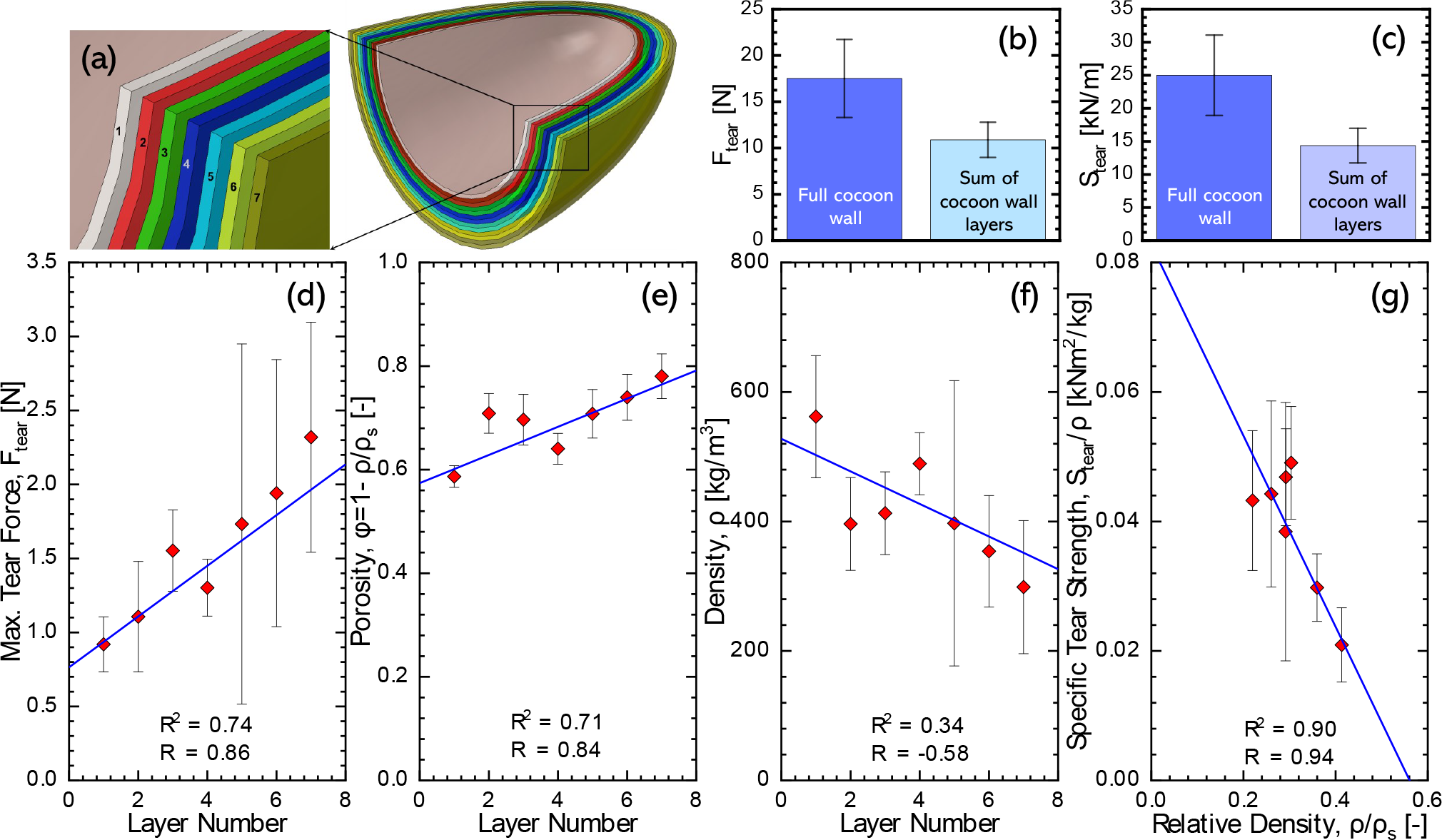
Tear statistics of layers from 1 (inner) to 7 (outer): (a) Layers of B. mori cocoon, (b) and (c) Comparison of tearing force and strength for the complete cocoon wall and the sum of the layers, (d) Tear force, (e) Porosity, (f) Density, and (g) Sp. tear strength against the relative density.

*F*_*tear*_ increases linearly from layers 1 to 7 (with a Pearson’s correlation coefficient (R) at 0.86, and a determination coefficient, (R^2^) of 0.74, Figure 4(d). We note that porosity (*ϕ*), also increases linearly from layers 1 to 7 starting at 0.59 in the innermost layer to 0.78 in the outermost layer (R = 0.84, R^2^ = 0.71), Figure 4(e). The density of cocoon layers naturally therefore decreases from layers 1 to 7 from 562 kg/m^3^ in the innermost layer to 299 kg/m^3^ in the outermost layer (R = -0.58, R^2^ = 0.34), as shown in Figure 4(f). Notably, the specific tear strength of the cocoon layers exhibited a strong negative correlation with relative density (R = 0.90, R^2^ = 0.94), Figure 4(g). Given that silk fibres constitute 74% of the cocoon mass, an increase in porosity from the innermost to outermost cocoon layers could be attributed to a reduction in the number of fibres per unit area (the areal density), which is in turn reflected by a decreasing density (cf. Figure 4(f)). The correlations observed in Figure 4 could be perceived as unusal, since cellular materials subjected to shear will typically show the reverse trends, in that their specific properties will rise, not fall, as a function of increasing relative density,^59^. Nevertheless, there is clear evidence from the literature that in fact torn textiles do behave in this manner^52,60–63^. This is because increased porosity in textiles provides additional spaces for fibre stretching, which distributes stress more evenly amongst neighbouring fibres. This resultantly reduces stress concentrations on individual fibres, or small clusters of fibres, thus enhancing the material resistance to tearing. This is schematised in Figure 5. Since each of these layers makes up the full cocoon wall, the mechanism for tearing described earlier (cf. Figure 3) will be comprised of both forms of tearing, allowing the *B. mori* cocoon have the composite properties of both stretch dominated failure, and brittle fracture.

**Figure 5:**
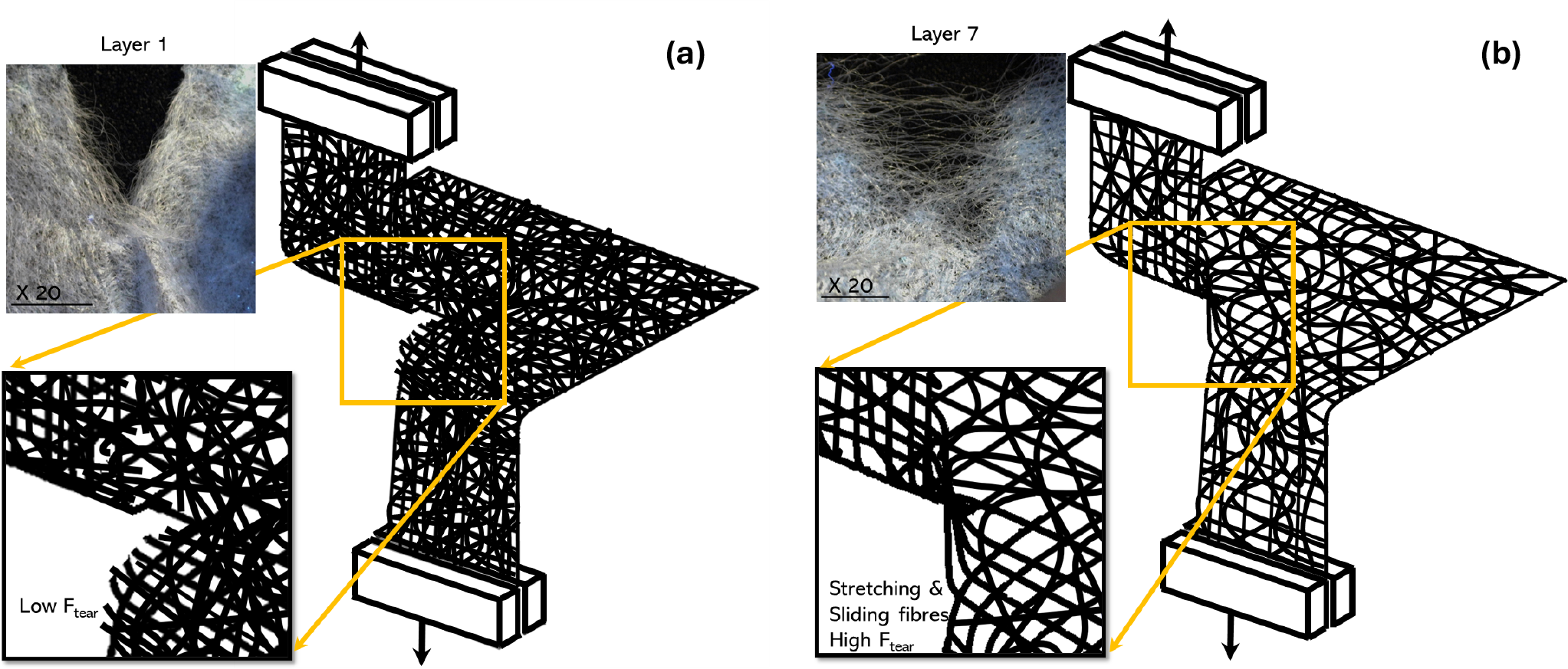
Tearing of individual *Bombyx mori* layers (a) low porosity leading to stress concentrations and low tear force (b) higher porosity leading to fibre sliding, stretching and high tear force. Insets: pictures of layer 1 and layer 7 respectively; scale bar = 1mm.

It can be be useful to know where the tear properties of *B*.*mori* cocoons fit within the broader spectrum of natural and engineering materials and textiles. Figure 6(a) shows an Ashby plot comparing *B. mori* tear energies and densities against a variety of materials including a range of textiles, elastomers, nonwovens, and films. A convex trade-off curve (grey line)^64^ is provided in Figure 6 where (1) identifies an optimal lightweight tear resistant material and (3) identifies the lower efficiency material that is both heavy and has low tear resistance. Trade off curves are useful as they elucidate the ‘good efficiency’ areas. In the case of Figure 6 that efficiency relates to balanced properties of density and tear resistance, which in this figure is identified as point (2) on the trade off curve. *B. mori* cocoon walls are therefore neither optimal lightweight tear resistant materials, nor are they inadequate as lightweight tear resistant materials. Similarly to many other natural materials, *B. mori* cocoon walls therefore exhibit a balanced trade-off between tear resistance and lightweightness^87,88^. An additional Ashby plot showing tear strength against density is provided as an Electronic Supplementary Figure 2. Figure 6(b) compares only the tearing energies of *B. mori* cocoon walls against a range of textiles (in both warp and weft), metals, polymers and films, elastomers, glass, hydrogel and cartilage. Glass has low values (0.026 kJ/m^2^) due to its inherent brittleness, while metals have very similar tearing energies to *B. mori* cocoon walls (50 kJ/m^2^). Textiles due to their interlaced architectures, dominate the histogram in terms of their resistance to tear with tougher high performance materials used in textiles, such as Kevlar and nylons exhibiting the highest tear energies. *B. mori* cocoon walls are natural non-woven architected fibrous materials and while their tear energies are not as high as systematically organised textiles (such as plain weave, twill weave, etc.) they still show respectable tear resistance, when compared against other natural materials such as cartilage (0.74 kJ/m^2^) .

**Figure 6:**
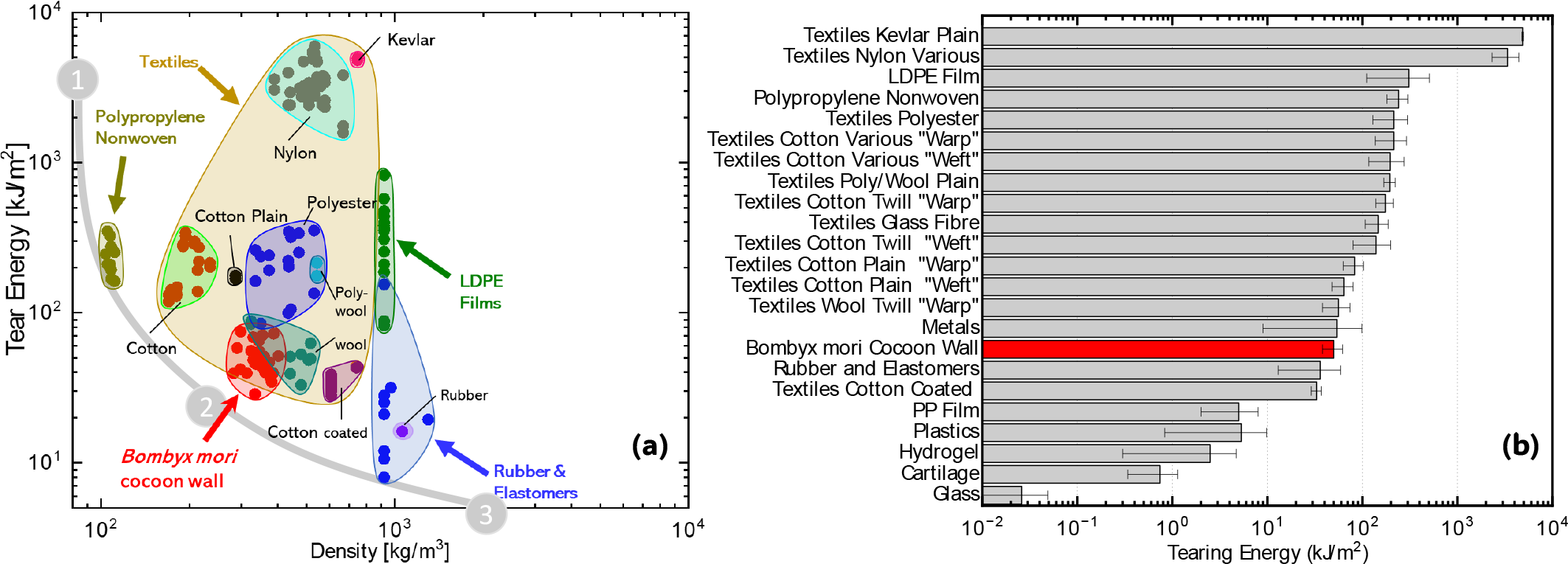
(a) Ashby chart showing the tearing energy of various materials (from^60–63, 65, 68, 69, 71, 73, 75, 80^) including *Bombyx mori* silk cocoon walls (this study) against density and (b) a histogram showing the tear energies (from^60, 62, 63, 65, 66, 68, 69, 71–86^) of a range of different materials/forms including the cocoon walls of *B. mori*. An additional Ashby plot showing tear strength (from^60–63, 65, 68, 69, 71, 73, 75, 80^) against density including *B. mori* silk cocoon walls (this study) is provided as: Electronic Supplementary Figure 2.

## CONCLUSIONS

*Bombyx mori* cocoon walls and its subdivided layers were tested in mode III tearing using trouser test methods. The cocoon wall requires 38% more force to tear than the sum of tearing loads of the seven subdivided layers and assuming an equal share of interface loading, each interlayer interface contributes 1.1 N of the total tear force. The tear force of 7 subdivided layers from the cocoon wall increased consecutively from the inner to the outer layers. Concurrently, the graded architecture of the cocoon wall also increased progressively in porosity and decreased progressively in areal density, from the inside to outside layer. These physical properties were found to be associated with an increased resistance to tearing forces. Inner layers are less porous and have very little space between the fibres, such that strain energy builds up and the fibres break in succession under an applied force. Fibres in the outer layers have a higher porosity with larger spaces and this allows for the stretching and sliding of individual fibres, leading to a distribution of strain energy over the neighbouring fibres, and consequently increasing the resistance of larger pore layers to tear force. A mechanism for the mode III tearing of full cocoon walls is suggested herein, where fibre sliding and stretching leads to fibre piling, followed by fibre breakage. The orientation of fibre piling determines the angle at which a cocoon wall will tear, and this orientation is presumably related to the presence of fibre junctions that act as nucleation sites for piling. The total failure of the cocoon wall comprising both higher and lower porosity layers, is therefore a composite exhibiting both brittle and stretch-dominated failure mechanisms.

## Supporting information

Comparison of standard tear force calculations

Electronic Supplementary Figure 1

Electronic Supplementary Figure 2

## DATA AVAILABILITY

Data for this publication will be made available through Edinburgh DataShare (https://datashare.ed.ac.uk/) and can also be made available from the corresponding author on request.

## ACKNOWLEDGMENTS

AUR wishes to thank the HEC Pakistan Scholarship made available through the National Textiles University, Pakistan.

## AUTHOR CONTRIBUTIONS

Conceptualization (PA); Data curation (AUR); Formal analysis (AUR, PA); Funding acquisition (AUR, PA); Investigation (AUR, VK, PA); Methodology (AUR, PA); Project administration (PA); Resources (PA); Software (AUR); Supervision (VK, PA); Validation (VK, PA); Visualisation (AUR, PA); Roles/Writing – original draft (AUR); Writing – review and editing (AUR, VK, PA).

## OPEN ACCESS STATEMENT

For the purpose of open access, the authors have applied a Creative Commons Attribution (CC BY) license to any author accepted manuscript version arising from this submission.

