## Supplementary material for "Mode III tear resistance of *Bombyx mori* silk cocoons": Comparison of standard tear force calculations

### Electronic Supplementary Material for:

#### Comparison of standard tear force calculations

Interpretation tearing varies by testing standard, since each standard recommends that tear force be calculated in different ways. This supplementary content compares the output from these standard test methods and bases calculations from tests conducted on 10 *Bombyx mori* cocoons. The following standards are cited in the original paper. The BS2782-3 method 360B uses the median of the plateau region of the tear force graph. BS ISO 34-1:2022 considers tear force as the median value from the middle 80% of the sample sets, disregarding, therefore, the upper and lower 10% values. BS EN ISO 6383-1:2015 and ASTM D1938-19 are similar in that they use the middle value to deduce the average tear force. The tear force in ASTM D2261 is the average of the five peak force values. BS EN ISO 13937-2:2000 determines the tear force by first dividing the force-extension graph into quartiles and eliminating values within the first quartile. From the remaining three quartiles, two greatest and two lowest peak values are then taken, and the tear force is determined by averaging these 12 measurements.

Figure A. below compares the tear force values produced for the same trouser tearing test using the various calculation techniques advised in testing standards. There was broad agreement in the interpretation of the tear force. The tear force found by measuring the single peak value of the force-extension graph (ASTM D624) and by averaging the five peak values of the graph (ASTM D2261) were found to be higher than the others, though they were in perfect agreement with one another.

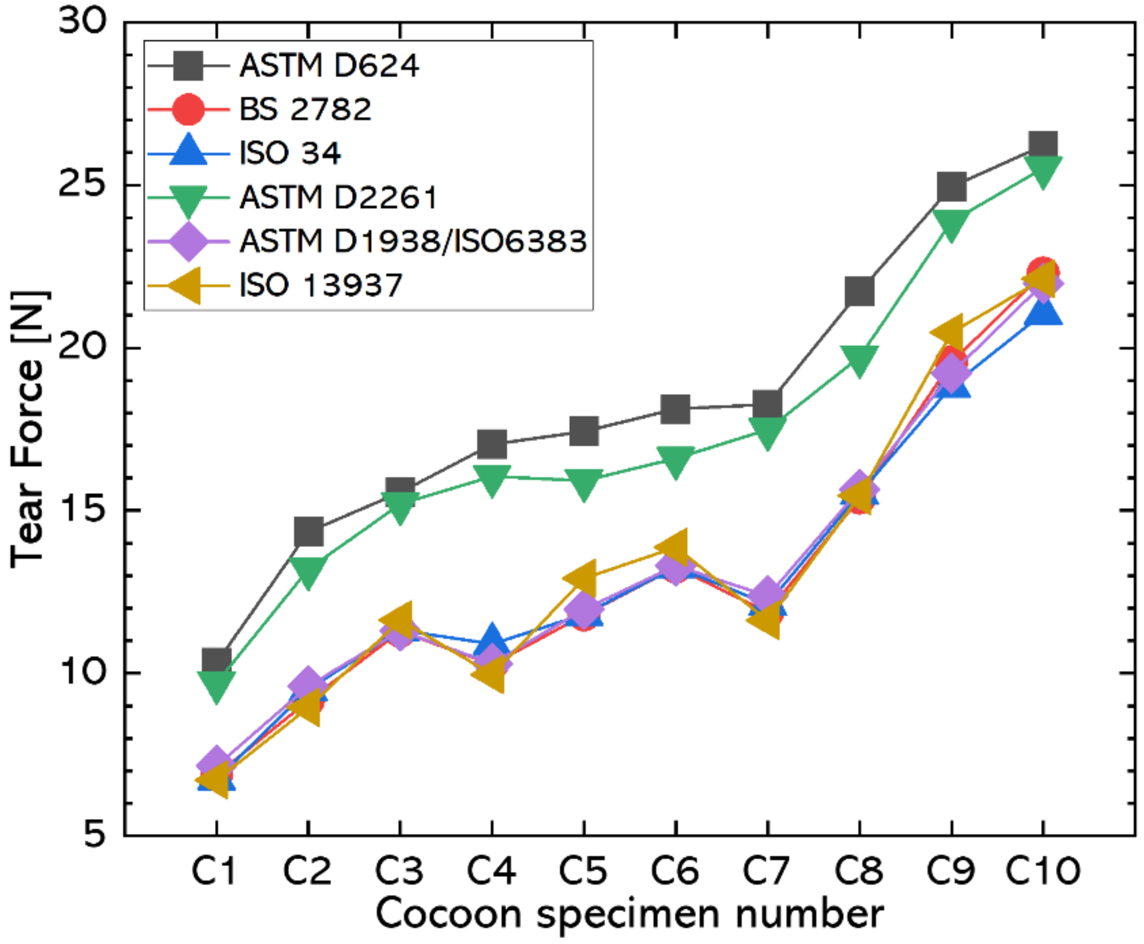


*Figure A. Comparison of tear forces calculated in accordance with different standards.*
