## Supplementary figures and images for "Mode III tear resistance of *Bombyx mori* silk cocoons"

### Electronic Supplementary Figure 1

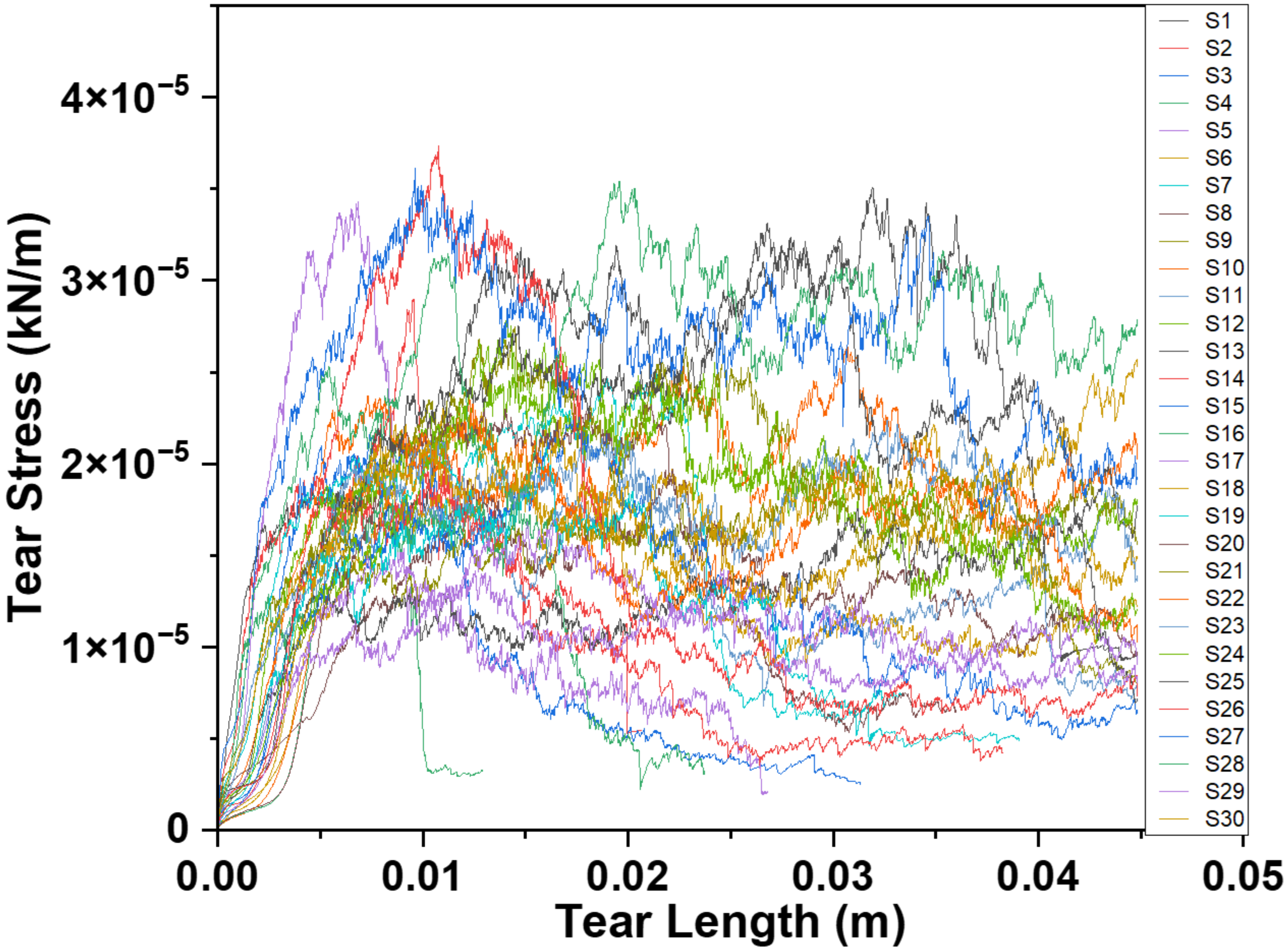


Tear stress vs. tear length for all *B. mori* cocoon tests (n = 30).
