## Supplementary material for "Mode III tear resistance of *Bombyx mori* silk cocoons": Electronic Supplementary Figure 2

Ashby plot showing tear strength vs. density for different materials and compared against *B. mori* silk cocoons.

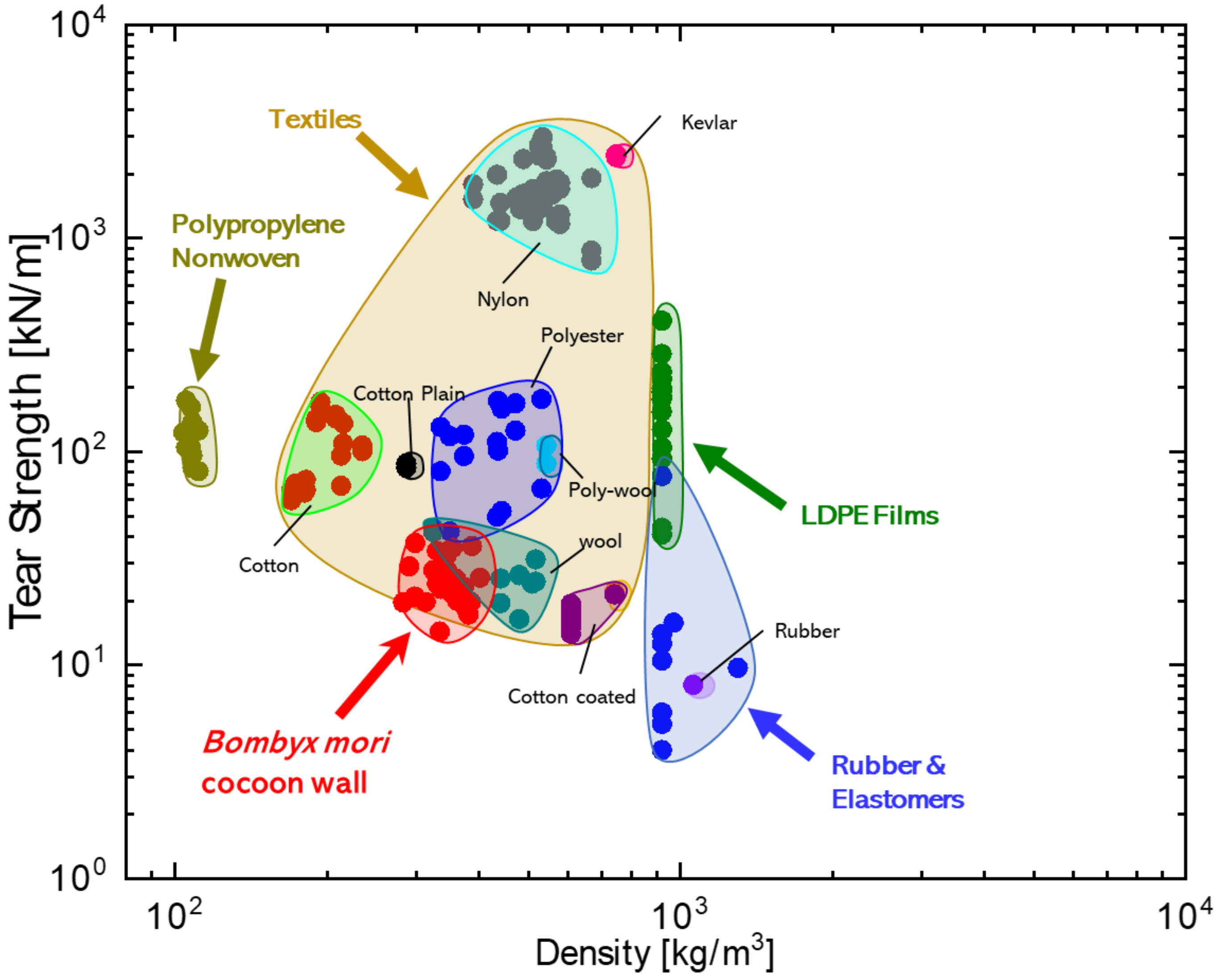
